# The Virtual Metabolic Human database: integrating human and gut microbiome metabolism with nutrition and disease

**DOI:** 10.1101/321331

**Authors:** Alberto Noronha, Jennifer Modamio, Yohan Jarosz, Nicolas Sompairac, German Preciat Gonzàlez, Anna Dröfn Daníelsdóttir, Max Krecke, Diane Merten, Hulda S. Haraldsdóttir, Almut Heinken, Laurent Heirendt, Stefanía Magnúsdóttir, Dmitry A. Ravcheev, Swagatika Sahoo, Piotr Gawron, Elisabeth Guerard, Lucia Fiscioni, Beatriz Garcia, Mabel Prendergast, Alberto Puente, Mariana Rodrigues, Akansha Roy, Mouss Rouquaya, Luca Wiltgen, Alise Alise Žagare, Elisabeth John, Maren Krueger, Inna Kuperstein, Andrei Zinovyev, Reinhard Schneider, Ronan M.T. Fleming, Ines Thiele

## Abstract

A multitude of factors contribute to complex diseases and can be measured with “omics” methods. Databases facilitate data interpretation for underlying mechanisms. Here, we describe the Virtual Metabolic Human (VMH, http://vmh.life) database encapsulating current knowledge of human metabolism within five interlinked resources “Human metabolism”, “Gut microbiome”, “Disease”, “Nutrition”, and “ReconMaps”. The VMH captures 5,180 unique metabolites, 17,730 unique reactions, 3,288 human genes, 255 Mendelian diseases, 818 microbes, 632,685 microbial genes, and 8,790 food items. The VMH’s unique features are i) the hosting the metabolic reconstructions of human and gut microbes amenable for metabolic modeling; ii) seven human metabolic maps for data visualization; iii) a nutrition designer; iv) a user-friendly webpage and application-programming interface to access its content; and v) user feedback option for community engagement. We demonstrate with four examples the VMH’s utility. The VMH represents a novel, interdisciplinary database for data interpretation and hypothesis generation to the biomedical community.

## Introduction

Metabolism plays a crucial role in human health and disease, and it is modulated by intrinsic (e.g., genetic) and extrinsic (e.g., diet and gut microbiota) factors. When considered individually, these factors do not sufficiently explain the development and progression of many complex non-communicable diseases, including metabolic syndrome and neurodegenerative diseases. Hence, a systems approach is necessary to elucidate the contribution of each of these factors and enable the development of efficient, novel treatment strategies.

Such a systems approach requires the easy sharing of knowledge and experimental data generated by different research communities. Databases represent a compelling method of storing, connecting, and making available a vast variety of information derived from primary literature, experimental data, genome annotations, etc. In fact, biological databases have become valuable tools for facilitating knowledge distribution and enabling research endeavors. The most recent release of the Nucleic Acid Research online Molecular Biology Database Collection contains more than 1,700 databases that span eight categories, including “metabolic and signaling pathways, enzymes and networks” and “human genomic variation, diseases, and drugs”^1^. Many of the databases span multiple categories, thus highlighting the increase in interdisciplinary research, which is of particular importance for systems biomedicine and synthetic biology^2^.

Despite the wealth of biochemical databases, a database that explicitly connects human metabolism with genetics, microbial metabolism, nutrition, and diseases has not been developed. One reason for the lack of such a database may be the use of non-standardized nomenclature, which complicates data integration. Moreover, manual curation of database content is time consuming and requires expert domain knowledge.

Genome-scale metabolic reconstructions represent the full repertoire of known metabolism occurring in a given organism and describe the underlying network of genes, proteins, and biochemical reactions^3^. High-quality reconstructions go through an intensive curation process that follows established protocols to ensure high standards and coverage of the information available on the organism^4^. Thus, metabolic reconstructions are valuable knowledge bases that summarize current information on metabolism within organisms. Genome-scale metabolic reconstructions have been generated for representatives of all domains of life, including humans^5^ and gut microbes^6^–^9^. Importantly, these metabolic reconstructions can be converted into computational models using condition-specific information, e.g., transcriptomic ^10^ or metabolomic data^11^,^12^. Open-access, community-developed toolboxes, such as the Constraint-Based Reconstruction and Analysis (COBRA) Toolbox^13^, facilitate simulations with metabolic models that permit us to address a variety of biomedical and biotechnological questions in silico^14^, ^15^.

Here, we describe the Virtual Metabolic Human (VMH, http://vmh.life) database, which consists of the five resources “Human metabolism”, “Gut microbiome”, “Disease”, “Nutrition”, and “ReconMaps”. These resources are interlinked based on shared nomenclature and database entries for metabolites, reactions, and genes (Figure 1). Overall, the VMH database contains 17,730 unique reactions, 5,180 unique metabolites, 3,288 human genes, and 632,685 microbial genes as well as 255 diseases, 818 microbes, and 8,790 food items. Unique features of the VMH database include i) metabolic reconstructions of human and gut microbes that can be used as a starting point for simulations; ii) seven comprehensive maps of human metabolism that permit a visualization of omics data and simulation results; iii) a nutrition designer that allows researchers to design personal dietary plans for computational simulations; iv) a user-friendly web interface for browsing, querying, and downloading the VMH database content; v) a well-documented representational state transfer application-programming interface (API) for easy access to the database content; and vi) user feedback through different platforms of the website, which will be curated and integrated into the knowledge base. Using four examples, we demonstrate how the resources in the VMH database can be exploited. Given the extensively curated, diverse information captured in the VMH database, this resource represents a unique, comprehensive, and multi-faceted overview of human metabolism.

**Figure 1:**
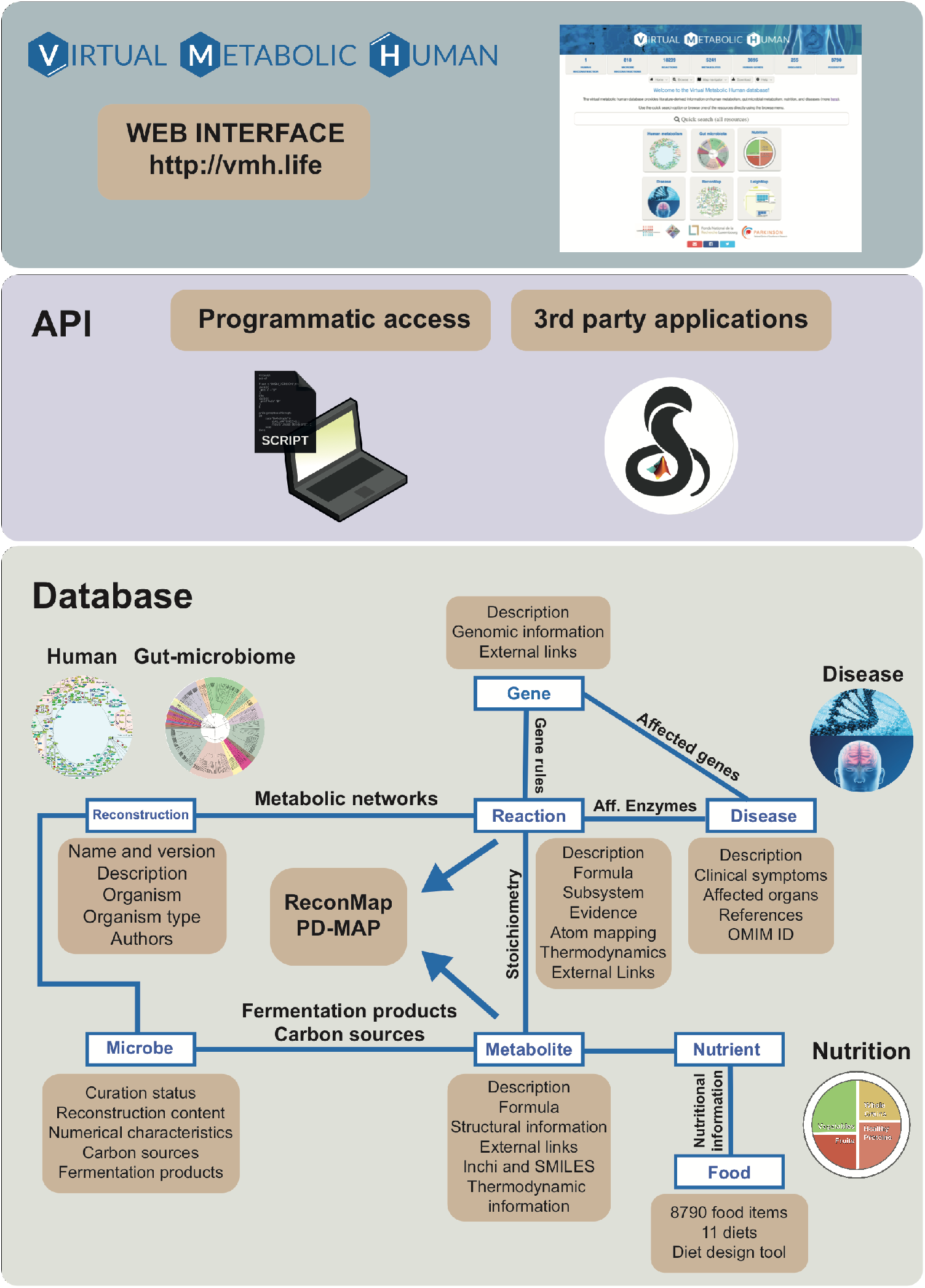
Overview of the Virtual Metabolic Human (VMH) database. The VMH database is divided into two interfaces, and its database contains five distinct but connected resources. Users can interact with the database using the two available interfaces: (i) a user-friendly web interface and (ii) an application-programming interface that allows programmatic access to the information contained in the database. At the core of the database is the representation of reconstructions as sets of reactions. The database connects the five resources through shared nomenclature: (i) the “Human metabolism” and “Gut microbiome” resources share metabolites and reactions, (ii) the nutrients in the “Nutrition” resource are mapped to metabolites that can be shared by the human and gut microbes, and (iii) the diseases in the “Disease” resource include affected genes and metabolite biomarkers present in the “Human metabolism” resource. Finally, the “ReconMaps” resource is connected to the “Human metabolism” resource via metabolites and reactions.

**Figure 2:**
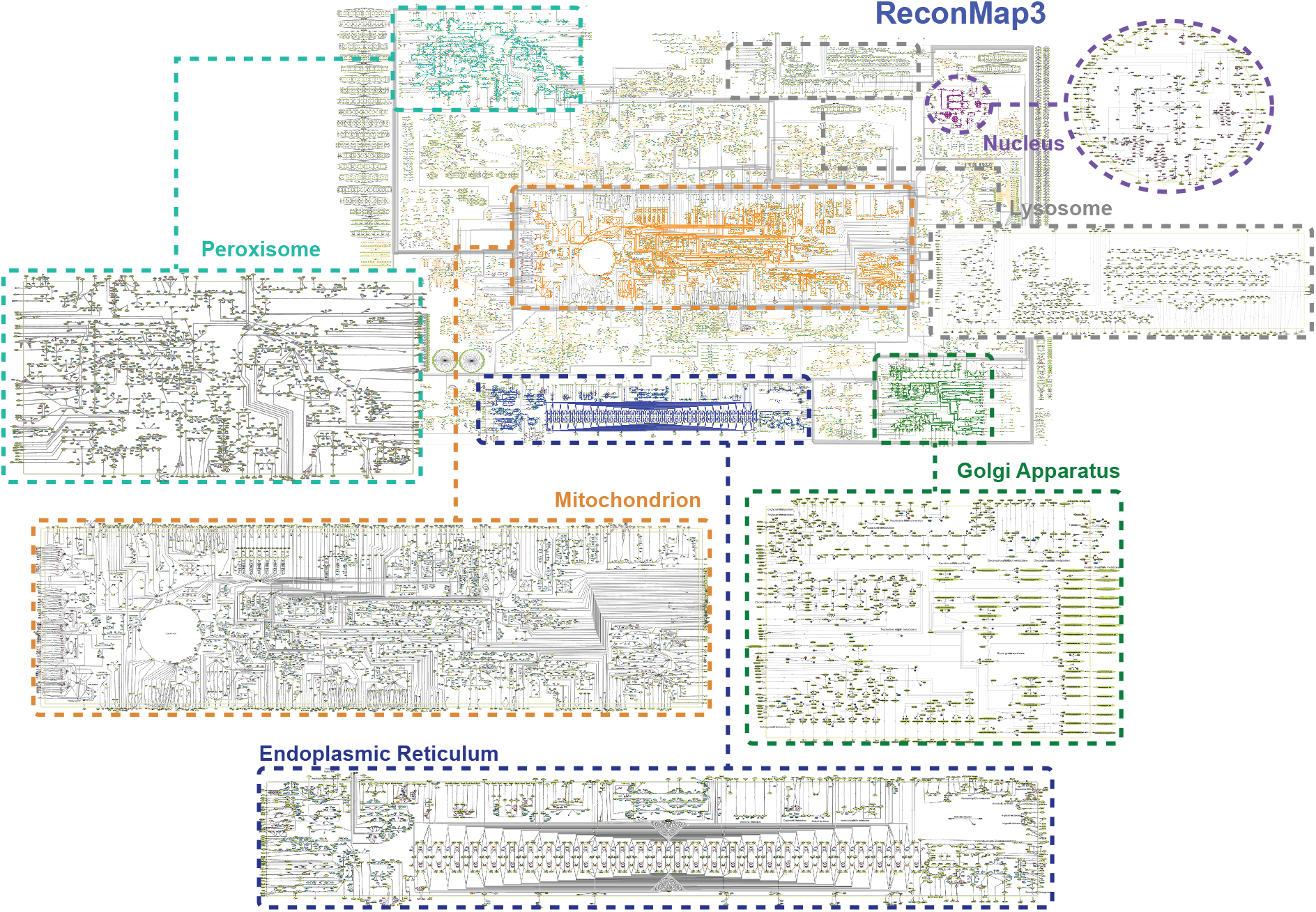
Visualization of the complexity of human metabolism with the “ReconMaps” resource. This resource consists of seven comprehensive maps for browsing of the content of the “Human metabolism” resource as well as for the visualization of experimental data and modeling results. While the ReconMap3 contains the metabolic pathways of all cellular pathways, the organelle-specific maps permit a more focused analysis of data and modeling results.

## Results

### “Human metabolism” resource

The VMH database hosts the most recent version of the human metabolic network reconstruction, Recon3D^5^, which describes the underlying network of 13,543 metabolic reactions distributed across 104 subsystems, 4,140 unique metabolites, and 3,288 genes expressed in at least one human cell. The content of Recon3D has been assembled through an extensive literature review over the past decade by the systems biology community^5^, ^16^–^18^.

Individual pages are dedicated to each reaction, metabolite, and gene. Such pages contain information on literature-based evidence as well the relations of the page subject to other entities in the VMH database (Figure 1). Novel features of Recon3D include molecular structures and atom mappings^5, 19^, which are visualized on the metabolite and reaction pages, respectively, in addition to thermodynamic information^20^. Great emphasis has been placed on collecting a comprehensive set of database-dependent and independent identifiers to allow for the identification of each entry and its cross-reference to more than 25 external resources (Table 1) that span metabolic, genetic, clinical, nutritional, and toxicological information sources.

### “ReconMaps” resource

The ReconMaps resource consists of seven human metabolic maps drawn manually using CellDesigner^21^ and hosted within the web service Molecular Interaction NEtwoRks VisuAlization (MINERVA)^22^. Six of these maps correspond to the cellular organelles found in human cells: the mitochondrion, nucleus, Golgi apparatus, endoplasmic reticulum, lysosome, and peroxisome. On the organelle level, reactions and pathways are drawn based on the defined subsystems, thus allowing the user to perform focused analyses of metabolism occurring in a particular cellular compartment. The seventh map, which is named ReconMap3, accounts for all six organelle maps plus the human metabolic reactions occurring in the cytosol and the extracellular space. Currently, ReconMap3 covers 8,151 of the 13,543 (60%) metabolic reactions and 2,763 of the 4,140 metabolites (67%) captured in Recon3D.

The maps support low-latency content queries and custom dataset visualizations, which are either represented as a text file or automatically uploaded from the COBRA Toolbox^13^, ^23^. Users can submit feedback through the map interfaces by right-clicking on specific elements. From each map entity, users can access the corresponding entry in the VMH database and obtain further information from external resources, such as the HMDB^24^, KEGG^25^, and CHEBI. Most notably, we connected the ReconMaps with the Parkinson’s disease map, PDMap^26^, which visualizes cellular processes known to be involved in Parkinson’s disease. The VMH, ReconMap3, and PDMap share 168 metabolites, thus providing a direct link between general human metabolism and Parkinson’s disease. Similarly, ReconMaps have been integrated via common players with the Atlas of Cancer signaling network resource (ACSN) containing mechanisms frequently deregulated in cancer. ACSN is a seamless network of 4600 biochemical reactions organizes into 52 functional modules covering 1821 proteins and 564 genes^27^. ReconMaps and ACSN share 252 proteins implicated in 22 functional modules of ACSN and in 51 subsystems of ReconMaps. This metabolic and cancer maps integration allows cross-talk studies and multi-omics data analysis in cancer^28^. Further disease maps are currently assembled by the community ^29^, and we will continue to increase the connectivity of the VMH and the ReconMaps to those valuable resources.

### “Gut microbiome” resource

The “Gut microbiome” resource contains 818 manually curated genome-scale metabolic reconstructions for microbes^6^ commonly found in the human gastrointestinal tract^30^ and belonging to 227 genera and 667 species. All microbial reconstructions were based on literature-derived experimental data and comparative genomics. A typical reconstruction contains an average of 774 (±275) genes, 1,218 (±249) reactions, and 944 (±143) metabolites. We provide detailed information for each strain and reconstruction along with known fermentation products and carbon sources, including supporting references. Importantly, this resource shares metabolite and reaction nomenclature with the other resources, thus allowing for an integrative analysis of microbial metabolism with host metabolism.

### “Disease” resource

The “Disease” resource links 255 inborn errors of metabolism (IEMs)^31^ to the genes present in the “Human metabolism” resource. A total of 288 unique genes and 1,872 unique VMH reactions are associated with these IEMs and provide biochemical and genetic descriptions. We have compiled clinical presentations, genotype-phenotype relationships, and the affected organ systems for each IEM from the primary and review literature. Additionally, we connect each entry with up to 12 external resources, thus providing further information on the diseases, genetic testing, and ongoing clinical trials.

The VMH database also hosts the Leigh Map^32^, which represents a computational gene-to-phenotype diagnosis support tool for mitochondrial disorders. The Leigh Map consists of 87 genes and 234 phenotypes expressed in Human Phenotypic Ontology (HPO) terms^33^, and they provide sufficient phenotypic and genetic variation to test the network’s diagnostic capability. The Leigh Map can be queried to generate a list of candidate genes and aims to support clinicians by providing faster and more accurate diagnoses for patients. This map will facilitate taking appropriate measures for further treatments and demonstrates the efficacy of computational support tools for mitochondrial disease.

### “Nutrition” resource

The “Nutrition” resource consists of two parts: i) a food database mapped onto the metabolites present in the VMH and ii) a diet database listing the nutritional composition of 11 pre-defined diets. The food database was built by integrating the molecular composition information for 8,790 food items distributed in 25 food groups obtained from the USDA National Nutrient Database for Standard Reference^34^. Of the 150 nutritional constituents, 100 could be mapped onto the metabolites present in the VMH database. Most of the remaining unmapped constituents represent general metabolite classes (e.g., fibers). The resource can be queried based on food items as well as their nutritional constituents.

The diet database contains 11 diets that were formulated based on real-life examples and literature. For instance, an “EU diet” was designed based on information from an Austrian survey ^35^. The diets consist of a one-day meal plan and include information on the energy content, fatty acids, amino acids, carbohydrates, dietary fibers, vitamins, minerals, and trace elements. The composition of each meal is given in appropriate portion sizes. The information for the nutritional composition of each food item and dish was obtained from the “Österreichische Nährwerttabelle” (http://www.oenwt.at/content/naehrwert-suche/). The molecular composition of a diet can be downloaded in grams per person (70 kg) per day or as a flux rate (in millimoles per person per day), which can be directly integrated with the human metabolic model ^5^, ^36^ using the COBRA Toolbox^13^.

### “Diet designer”

The “Diet designer” tool allows users to design their diets. The interface is divided into two lists: “Available foods” and “Selected foods”. Users can search and select any of the available 8,790 foods and add them to the list of selected foods by specifying a portion size. As the user designs the diet, the overall information is updated for total calories, lipids, proteins, carbohydrates, and weight. The user can then see and download the corresponding molecular composition and flux values for tasks that include computational modeling.

Overall, the VMH knowledge base combines four well-curated resources on human metabolism in a unique and novel manner that will aid researchers in their biomedical applications.

### VMH database accelerates the generation of novel hypotheses

We demonstrate four possible applications that exploit the versatility of the VMH database, its multiple resources, and the facilitated access to other knowledge bases through the provided external links. Many complex queries can be performed through the VMH database’s comprehensive API, and the obtained data can be analyzed using third-party tools.

### Exploring the complex interactions between microbes, nutrition, and host metabolism

Microbial metabolic interactions represent driving forces for the microbial community composition as well as emergent metabolic properties, e.g., the production of short-chain fatty acids, which serve as an energy source for the human body and have an immunomodulating role^37^. By using the VMH database, we can systematically query for shared metabolite exchanges among humans, microbes, and nutritional resources. The portfolio of exchange reactions, which define metabolites that can be exchanged between an organism and its environment, provides an organism-specific “interaction profile”, and a comparison of these profiles provides a better understanding of the roles that specific organisms can play in complex systems, such as the human gut. All 818 gut microbes share a common set of 17 exchange reactions, whereas each microbe has an average of 132 ± 27 exchange reactions. We compared the presence of exchange reactions across these 818 gut microbes, using tSNE for visualization of the high-dimensionality data^38^. We found a separate cluster (Figure 3-A) formed mainly by 110 representatives belonging to the phylum Bacteroidetes, which indicated that these microbes share a unique set of exchange reactions distinct from most microbes of the other phyla. In contrast, the other phyla overlapped more in their exchange reactions and thus in their interaction potential. We then compared the pan-metabolic repertoire between Bacteroidetes and Firmicutes, i.e., all metabolites and reactions found in any of their representatives. As expected, most of the repertoire was shared between the two phyla (Figure 3-B). Of the 165 metabolites specific to Firmicutes, 42 had corresponding exchange reactions, whereas of the 61 metabolites specific to Bacteroidetes, 48 had corresponding exchange reactions. In the VMH database, information on the pathways, including for these specific metabolites, can be retrieved via the “Subsystem” attribute of the corresponding reactions. The Firmicutes-specific exchanged metabolites are involved in 66 subsystems, which are mostly associated with the metabolism of amino acids, whereas the Bacteroidetes-specific exchanged metabolites display an ability to degrade components of plant cell walls, such as polysaccharides and proteoglycans. Additionally, compared with other phyla, Bacteroidetes interact less with the nutrition resource (Figure 3-C), indicating that the members of this phylum rely on the availability of a smaller variety of nutrients. Bacteroidetes representatives, e.g., *Bacteroides* and *Prevotella* spp., efficiently use host glycans and/or diet-derived fibers^39^, which has been accounted for in the AGORA resource through extensive literature-based curation^6^. Plant polysaccharide and proteoglycan contents of foodstuffs are currently not reported in nutritional databases, such as the USDA^34^, thus explaining the low overlap of fibers and glycans usable by the Bacteroidetes from the Nutrition resource. Recent efforts have focused on closing this gap by metabolomically characterizing foodstuffs, e.g., FoodDB (http://foodb.ca/). Overall, we found that the gut microbes overlapped on average with 41 ± 5 of the 102 defined food components, and these overlapping metabolites mostly belonged to the subsystems (excluding transports) fatty acid oxidation, cholesterol metabolism, and fatty acid synthesis (Figure 3-C). For the overlap with the “Human metabolism” resource, we found an average of 94 ± 17 interactions, which were mainly in nucleotide interconversion, glycerophospholipid metabolism, and methionine and cysteine metabolism (Figure 3-D), indicating that these metabolites could potentially be exchanged between gut microbes and humans. Hence, the VMH database enables a microbiota-wide systematic approach to exploiting and characterizing the capabilities of the gut microbiota to metabolize dietary items and thereby modulate host metabolism.

**Figure 3:**
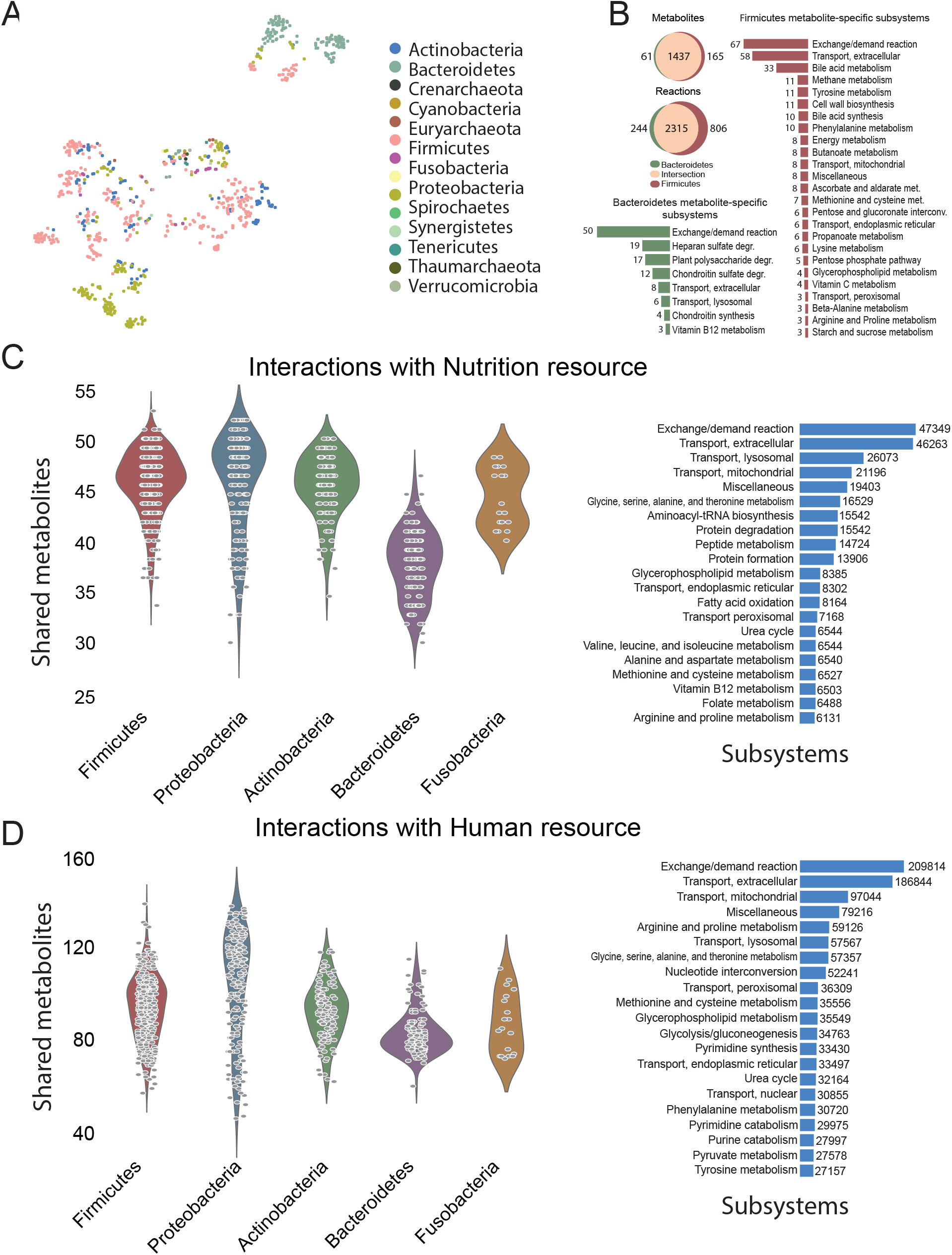
“Gut microbiome” resource enables the comparison and analysis of gut microbial metabolic capabilities. **A**. tSNE of the interaction profile of the microbes in the VMH database. Only exchange reactions were considered because they represent potential interactions. **B**. Reaction and metabolite content comparison of the two most abundant phyla in the VMH database: *Bacteroidetes* and *Firmicutes*. **C**. Comparison of interactions between phyla and the “Human metabolism” resource and the “Nutrition” resource.

### Designing synthetic microbial communities with the VMH database

Synthetic microbial communities, which capture key properties of more complex communities, are commonly used to test biomedically relevant hypotheses and gain novel insights into microbial ecology^40^, ^41^. For instance, Desai et al.^40^ have demonstrated that in a 13-species synthetic gut microbiota, four species were capable of growing on mucus-O-glycans. The exchange profile of the corresponding four microbial metabolic reconstructions was in with the experimental data for almost all glycans (Figure 4-A). Only the metabolic reconstruction of *Barnesiella intestinihominis* fails to capture some of the experimental observations^40^ since this information was unavailable at the time of the curation of AGORA. This comparison highlights the importance of the manual, literature-based curation effort undertaken for our microbial reconstructions and the iterative nature of reconstruction refinement^4^ due to the increasing availability of new experimental data.

**Figure 4:**
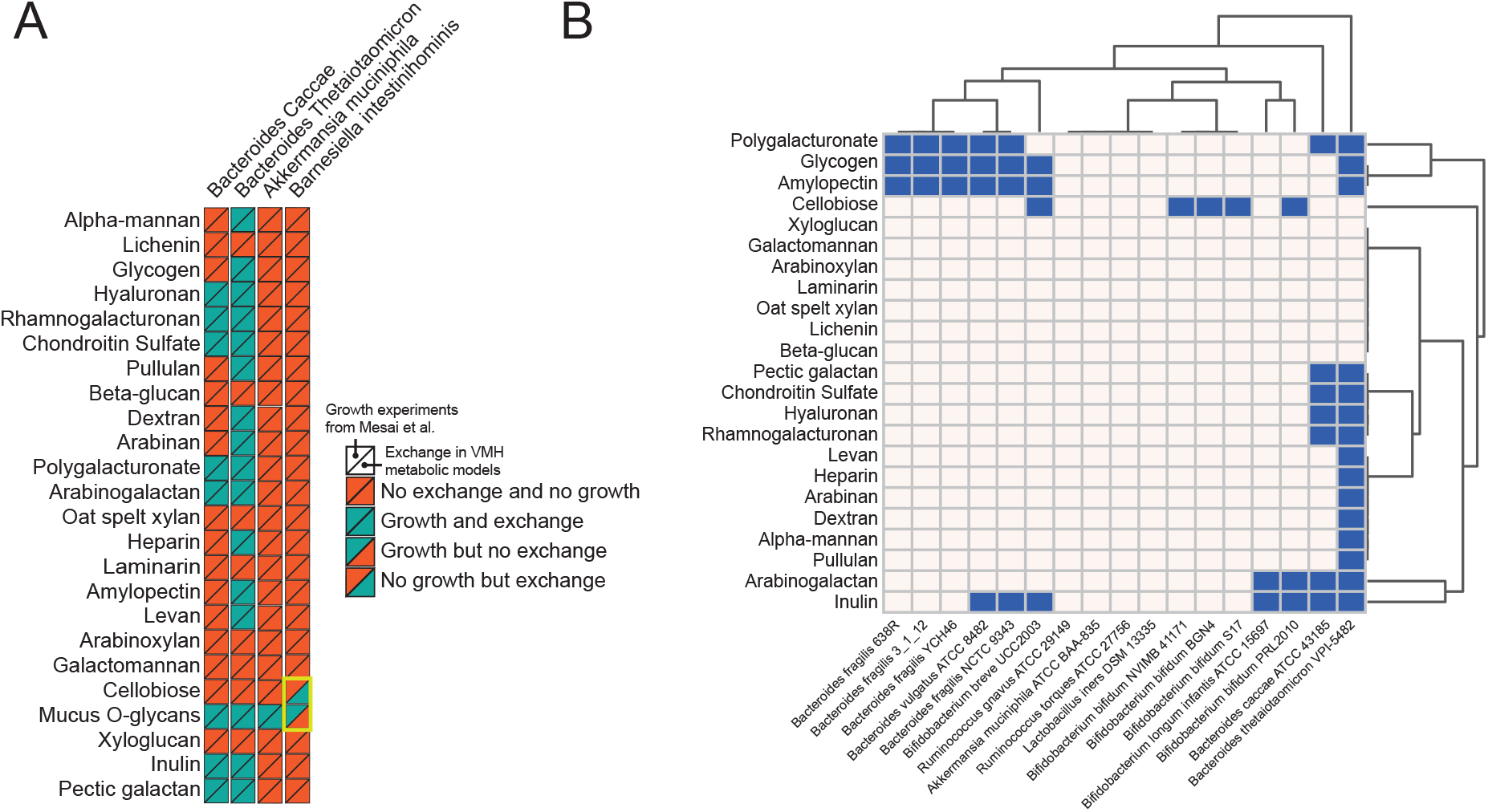
Experimental design of synthetic microbiotas using the VMH database. **A**. Comparison between AGORA models and experimental results. The existence of an exchange reaction (ability to uptake a given compound) was compared against single-carbon-source growth experiments (Desai et al., 2016). Full concordance was found except with *Barnesiella intestinihominis*. **B**. Other microbes in the VMH database displaying the ability to take up mucus O-glycans demonstrate how the VMH database can be used for the design of synthetic microbiotas.

We used the VMH database to design synthetic gut microbial communities with a particular glycan-degradation profile. To this end, we identified 14 gut microbes that mostly belonged to *Bacteroides* and *Bifidobacterium*, and an overlapping mucus-O-glycans utilization profile was used for the four microbes (Figure 4-B). From the VMH database, we can retrieve an extended glycan and polysaccharide utilization profile, which could be used to broaden the carbon source utilization capabilities of the synthetic microbiota. This example illustrates that the VMH database enables the user to analyze the *in silico* potential of different microbes and supports the design of experiments by exploiting the collection of literature-based and manually curated subsystems, i.e., “Fermentation products” and “Carbon sources”, available in the “Gut microbiome” resource.

### Drug detoxification and retoxification

Xenobiotic metabolism often involves glucuronidation of drugs, which is an important mechanism for drug detoxification and subsequent elimination through bile or urine^42^. UDP-glucuronic acid (VMH ID: udpglcur) is formed in the liver and represents an essential intermediate in this process (Figure 5-A), and its availability can be rate-limiting for the elimination of exogenous and endogenous toxins in rats^43^. Within the VMH, 18 genes encode enzymes that carry out the liver glucuronidation of 37 endogenous and exogenous metabolites, including 18 drug metabolites^5^, ^44^.

**Figure 5:**
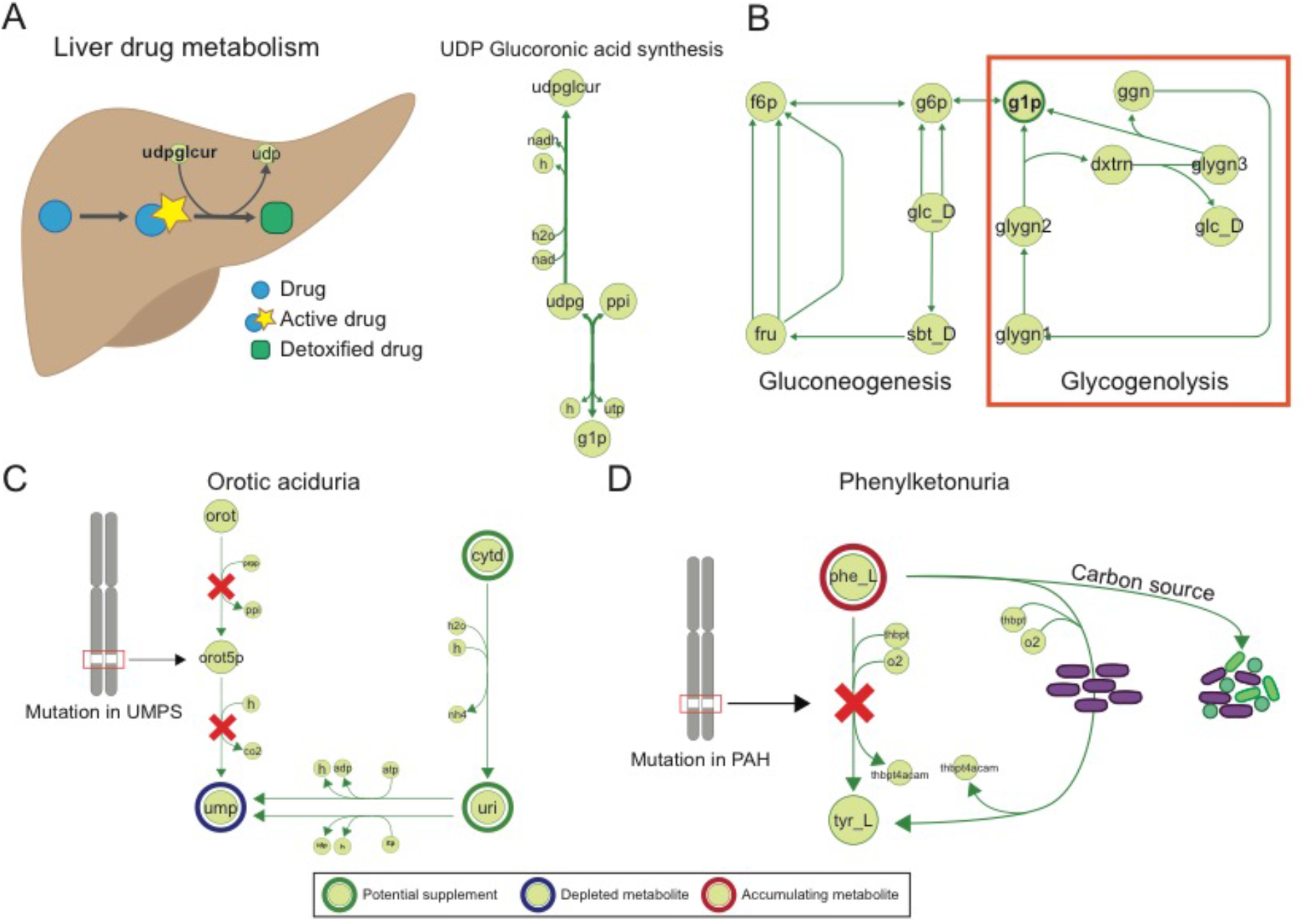
Using the VMH database to investigate mechanisms of disease and drug metabolism. **A**. UDP-glucuronic acid (VMH metabolite: udpglcur) is formed in the liver and an essential intermediate in the glucuronidation of drugs. UDP-glucose 6-dehydrogenase (VMH reaction: UDPGD) converts UDP-glucose (VMH metabolite: udpg) to UDP-glucuronic acid, and UTP-glucose-1-phosphate uridylyltransferase (VMH reaction: GALU) converts glucose-1-phosphate (g1p) to UPD-glucose. **B**. Sources of g1p found in ReconMap3. In rats, glycogenolysis is the source of UDP-glucuronic acid in the process of glucuronidation^45^. **C**. Orotic aciduria: mutation in UMPS affects reactions that transform orotic acid into uridine monophosphate. VMH pathway information points to known treatment with the use of uridine and cytidine. **D**. Phenylketonuria: mutation of PAH causes an inability to degrade phenylalanine. Certain gut microbes show the ability to degrade phenylalanine or use it as carbon source. A treatment strategy might involve gut microbiome community engineering.

To identify potential dietary intervention strategies to alleviate UDP-glucuronic acid limitation, we used the VMH database to investigate UDP-glucuronic synthesis pathways. Briefly, UDP-glucuronic acid is synthesized from UDP-glucose (VMH: udpg) by the reaction of UDP-glucose 6-dehydrogenase (VMH ID: UDPGD), which in turn is synthesized by UTP-glucose-1-phosphate uridylyltransferase (VMH reaction: GALU) from glucose-1-phosphate (VMH ID: g1p) and UDP-glucose (Figure 5-A). Glucose-1-phosphate in turn is generated by the pathways “Gluconeogenesis” and “Glycogenolysis” (Figure 5-B). At least in rats, the UDP-glucuronic acid used for glucuronidation is predominantly derived from glycogen^45^. Accordingly, a high dosage of acetaminophen can deplete the liver glycogen storage^43^. The importance of gluconeogenesis for glycogen storage is highlighted by the fact that two out of 14 Mendelian glycogen storage diseases listed in the VMH database are due to defects in the enzymes of this pathway. Additionally, liver glycogen storage can be effectively replenished by carbohydrates, such as glucose and fructose, after exercise^46^. Interestingly, maltodextrin (MD) drinks containing galactose or fructose are twice as effective at restoring post-exercise liver glycogen synthesis as MD drinks rich in glucose^47^. However, maltodextrin has a higher glycemic index than sugar, and it can impair intestinal anti-bacterial responses and defense mechanisms^48^ as shown by the increased survivability of Salmonella^49^. Since the absorption of fructose is facilitated when ingested in combination with glucose^50^, we searched the VMH database for foodstuffs that are high in fructose and glucose (Supp. Table 1). Naturally occurring foodstuffs include honey, medjool dates, and raisins (Supp. Table 1-A). The content of galactose is relatively low in most food items, although honey and Greek yogurt are among the best choices (Supp. Table 1-B). Thus, naturally occurring foodstuffs may be used to replenish glycogen stores, thereby providing the necessary glycogen precursors for gluconeogenesis.

Once glucuronidated, the drug derivatives are excreted either via urine or the enterohepatic route. In the latter case, the glucuronidated drug, such as the cancer drug irinotecan, can be retoxified through the action of microbial beta-glucuronidase^51^, ^52^. Five AGORA microbes have been demonstrated experimentally to possess the beta-glucuronidase enzyme and thus use glycans containing glucuronate as carbon sources. This information is captured by the VMH resource (corresponding VMH reactions: GLCAASE8e, GLCAASE9e, GLCAASE_HSe, GLCAASEe). This example demonstrates how the VMH database can provide a novel, multi-faceted view of human drug metabolism based on a consideration of dietary and microbial aspects.

### Probiotic approaches to rare disease treatment

Orotic aciduria (OMIM 258900) is an autosomal recessive disorder caused by a mutation in the uridine monophosphate synthetase gene (EntrezGene ID: 7372). In the VMH database, this gene is associated with two reactions (VMH ID: ORPT and OMPDC) that transform orotic acid (VMH ID: orot) into uridine monophosphate (VMH ID: ump; Figure 5-C), which is consistent with the two enzymatic activities encoded by this gene^53^. The gene deficiency leads to pyrimidine starvation, which can be effectively treated with uridine or cytidine (Figure 5-C). However, the supplemented uridine competes for intestinal absorption with dietary pyrimidines or purines^54^,^55^. The VMH accounts for the corresponding facilitated transport reactions associated with the *SLC29A1* (EntrezGene ID: 2030) and *SLC29A2* (EntrezGene ID: 3177) as well as the sodium-dependent transport reaction enabled by *SLC28A3* (EntrezGene ID: 64078). We have previously predicted that the human commensal gut microbe *Bacteroides thetaiotaomicron* could also supplement the host with uridine^56^. By using the VMH database, we can identify 438 additional gut microbes that could potentially supplement the human host with uridine because they encode the 5’-nucleotidase (VMH ID: NTD2, E.C. 3.1.3.5) as well as a uridine transporter (VMH ID: URIt2r). Of those microbes, 18 have been classified as probiotics in the VMH database and include 15 *Bifidobacterium* strains, two *Clostridium butyricum* strains, and *Lactobacillus reuteri* MM4–1A. These probiotics are commonly found in yogurts, fermented food products, and probiotic formulations. Although we could not find evidence for probiotic use in orotic aciduria, guidelines for the management of methylmalonic and propionic acidemia included the use of probiotics^57^. Furthermore, researchers have demonstrated the benefit of engineered *L. reuteri* strains in a murine phenylketonuria (PKU) model^58^.

PKU is caused by a mutation in the gene PAH (EntrezGene ID: 5053), and it leads to an inability to degrade phenylalanine (Figure 5-D). The life-long treatment consists of a diet low in phenylalanine. An alternative strategy could be to engineer the gut microbiota such that it consumes the excess of this compound. One option is the aforementioned engineering of probiotics, where the researchers introduced the phenylalanine ammonia-lyase (EC 4.3.1.24) to L. reuteri. This enzyme is ubiquitous in higher plants but rare in microbes, and the VMH database does not account for the corresponding reaction (although that does not mean that none of the 818 microbial genomes encode this gene). However, an alternative pathway (VMH IDs: PHETA1, PLACOR, PLACD) that converts phenylalanine to trans-cinnamic acid occurs in six strains of *Clostridium* (*Clostridium difficile* NAP08, *Clostridium difficile* NAP07, *Clostridium difficile* R20291, *Clostridium sporogenes* ATCC, *Clostridium difficile*, and *Clostridium sticklandii* DSM). Another option may be to “replace” the mutated PAH gene with a microbial counterpart. In the VMH database, 26 microbes encode the microbial version of the genes, including two commensal *Bacillus cereus* strains and one probiotic strain (*L. reuteri* SD2112). Although *B. cereus* is known to be a causative agent in a minority of foodborne illnesses^59^, the *L. reuteri* strain has been added to yogurt formulations with the aim of improving oral hygiene^60^. Additionally, in the VMH database, the literature-based table for carbon sources lists three additional commensal microbes that use phenylalanine as a carbon source: *Clostridium bartletti DSM 16795*, *Anaerobaculum hydrogeniformans* ATCC BAA-1850, and *Gordonibacter pamelaeae* 7-10-1-bT. The latter two have been recently patented to be used as probiotics for the inhibition of clostridial-caused inflammation^61^.

Taken together, the VMH database can be used to identify candidate microbes that could be used in addition to or as a replacement for current dietary intervention strategies used in the treatment of inborn errors of metabolism.

## Discussion

The VMH database captures information on human and gut microbial metabolism and links this information to hundreds of diseases and nutritional data. Therefore, the VMH database addresses an increasing need to facilitate rapid analyses and interpretations of complex data arising from large-scale biomedical studies. The four sample applications illustrate how the VMH resources enable broad-type analyses that would otherwise require diverse expertise and the use of numerous databases and textbooks. While the front-end of the VMH database permits complex, interdisciplinary queries by the general user, the comprehensive API enables programmers to perform many complex searches on the database content.

An integral part of systems biology is computational modeling. Constraint-based modeling is gaining increased attention by a broad interdisciplinary scientific community. At the foundation of the VMH database lies genome-scale metabolic reconstructions (Figure 1, 6). Thus, the VMH database directly serves the growing constraint-based modeling community and its needs by providing a user-friendly interface for reconstructing content. These reconstructions are provided in multiple standard formats (e.g., SBML) for download and allow for access to the entire knowledge base via the API and enable the *in silico* formulation of personalized diets via the “Diet designer”, which can then directly be integrated with the human or microbial metabolic reconstructions using the COBRA Toolbox^13^. Simulation results based on these diets^36^,^62^,^63^ or based on the integration of omics data, e.g., metabolomics^12^ and transcriptomics^64^, can then be visualized and interpreted in the context of human metabolic and disease-specific maps, thus providing a more comprehensive, systems-level understanding of the modeled system.

**Figure 6:**
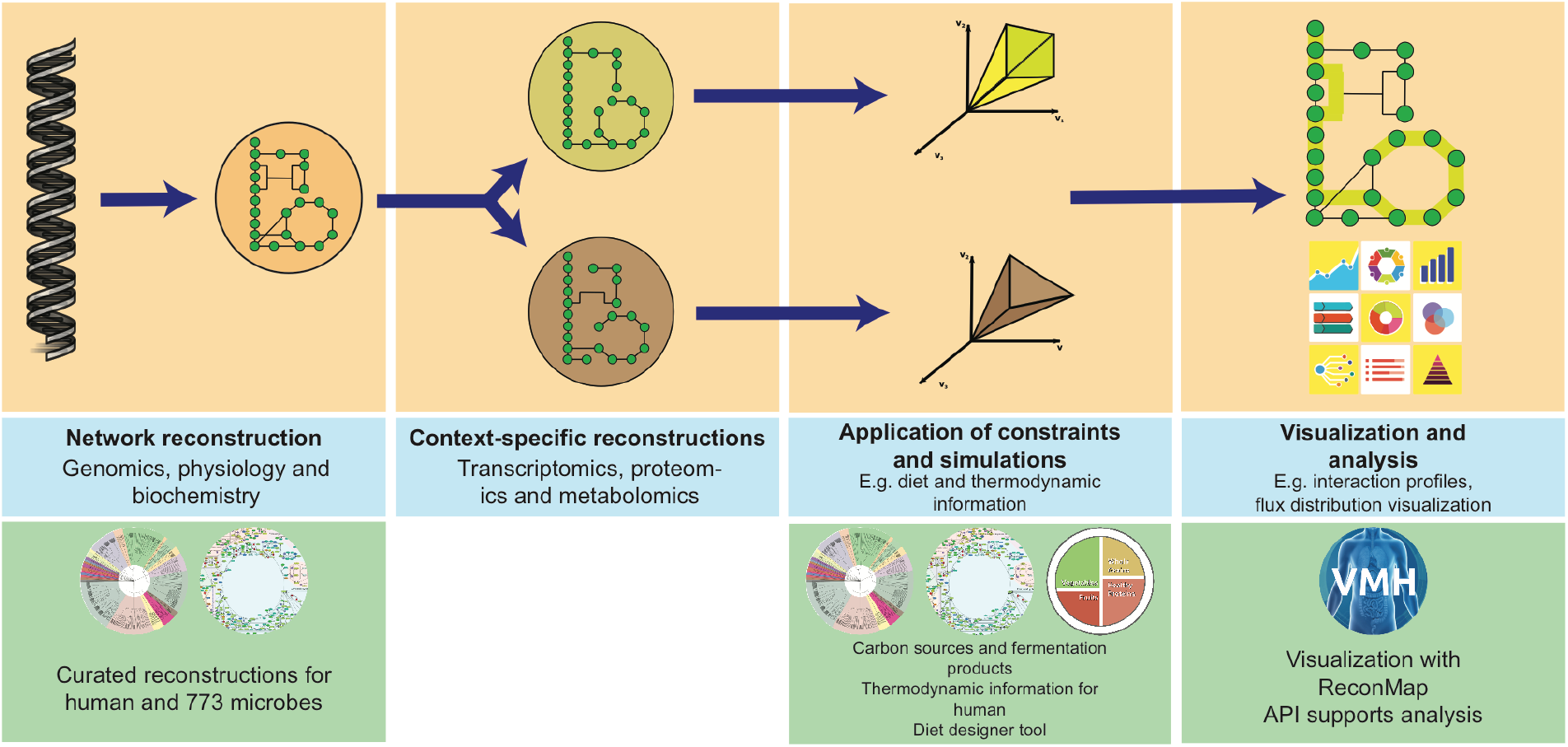
Overview of the Constraint-Based Reconstruction and Analysis (COBRA) methodology. From the genome annotation of an organism, a general reconstruction is built, and with the integration of data from different sources, context-specific reconstructions can be derived. Adding constraints, such as media for microbial organisms or diet for humans, allows for the derivation of a model that can perform simulations to predict the behavior of the cell/organism under different conditions. The VMH database supports this pipeline with its resources on various levels, from providing a set of curated reconstructions to information that can be included as constraints to the visualization of simulation and experimental results with ReconMaps.

A growing number of studies link microbial composition with diet and disease^65, 66^. The generation of novel hypotheses about the functional implications of observed correlations, e.g., between microbial abundances in disease states, is hindered by the lack of online databases to facilitate such work. In particular, the “diet designer” tool in conjunction with computational modeling permits the generation of *in silico* hypotheses that could then be experimentally tested. The use of synthetic microbial communities is of great value for hypothesis testing, and the VMH database can facilitate the design of defined microbial communities with specified metabolic capabilities.

Unique and distinguishing features of the VMH database are the three key factors. First, the VMH database is a comprehensive, interdisciplinary database that permits complex queries. Second, the VMH database provides a graphical representation of the “Human metabolism” resource through the “ReconMaps” resource, thus allowing for the analysis of complex, multi-faceted omics data in the context of the biochemical knowledge captured in the VMH database. Third, the VMH database represents a starting point for computational modeling of human and microbial metabolism in healthy and diseased states by providing information and simulation constraints and being fully compatible with the COBRA Toolbox^13^. As such, the VMH database provides a novel research tool by increasing the availability of diverse data along the diet-gut-health axis to the biomedical community.

## Acknowledgments

This study received funding from the Luxembourg National Research Fund (FNR) through the ATTRACT program (FNR/A12/01), the CORE program (C16/BM/11332722), the OPEN grant (FNR/O16/11402054), the National Centre of Excellence in Research (NCER) on Parkinson’s disease, the European Union’s Horizon 2020 research and innovation program under grant agreement No. 668738 and the European Research Council (ERC) under the European Union’s Horizon 2020 research and innovation program (grant agreement No 757922).

## Methods

### Technical implementation

The VMH database was implemented with MySQL 5.6 (https://dev.mysql.com/). A relational-database model was chosen for its simplicity and flexibility. The front-end is reachable via web browser at http://vmh.life and was developed in Sencha ExtJS 5.1 (https://www.sencha.com/). The API was developed using Python 2.7 with the DJANGO framework and the Django Rest Framework package. The API can be used by third-party applications and is also accessible via web browser at https://vmh.uni.lu/_api/ and its documentation page at https://vmh.uni.lu/_api/docs.

### Detailed pages

The VMH database contains detailed information for each entity in the database as well as numerous external and internal links. Through the user interface, a user can search the different resources and navigate the various levels of detail, e.g., from disease information to low-level metabolite biomarker information and chemical structure.

### Metabolite page

Each metabolite in the VMH database is represented by an abbreviation that uniquely identifies a specific molecule involved in at least one metabolic reaction present in the database. Each metabolite also contains a name that better identifies that specific molecule, and the description and synonyms are extracted from HMDB^2^4 when available.

The charged metabolite formula represents the acid/base form of the neutral molecule (at pH 7.2), and the associated charge values are provided. Inchi strings^67^ and Smiles^68^ are available for most of the metabolites in the VMH database^19^. Mol files were generated for most Recon3D metabolites, and with the use of ChemDoodle, users can generate similar structural visualizations (and interactions) of these metabolites as provided by many other chemical databases. The mol files are also available for download on the metabolite page. Thermodynamic information is provided when available^20^ along with details on how it was calculated. The standard Gibbs energy is displayed for the different cellular compartments. Information on the presence of metabolites in human biofluids was obtained from HMDB^24^, literature sources^69^–^74^, and the Netherlands Metabolomics Centre (NMC – http://www.metabolomicscentre.nl/). Biofluid information can be qualitative (presence/absence) or quantitative if the range of values is specified. The sources of the information are specified in each row of values. Biomarker information connects the metabolite with the disease resource^31^.

Each metabolite includes information on the number of human and microbial reactions it is involved in as well as whether it is a carbon source or a fermentation product of any of the 818 microbes.

### Reaction page

A VMH reaction is represented by its abbreviation and a detailed description, which includes the name of the enzyme that catalyzes the reaction and the cellular compartment in which the reactions occur. For instance, to represent the transport of D-3-amino-isobutyrate, the VMH entry is written as D_3AIBt: D-3-Amino-Isobutyrate Transport Formula: 3aib_D[c] –> 3aib_D[e] which represents the transport from cytosol to extracellular environment. Additionally, we provide information on the associated subsystem, notes added by the model curators, a confidence score^4^, and the literature sources. Each reaction is graphically displayed in an atom-mapped fashion, and its structure is also available through a view that uses the software ChemDoodle. KEGG^25^, ReconMaps, and COG^75^ identifiers along with the Enzyme Commission number are displayed under “External Links”. The standard reaction Gibbs energy is displayed when available. Finally, from the reaction page, a user can navigate to associated genes, microbes, and diseases.

### Human gene page

The gene number used in the VMH database to identify human genes is a combination of Entrez Gene identifier and the transcript ID (specified with ‘.1’, ‘.2’, or ‘.3’ after the Entrez Gene ID). The use of transcripts explains why the genes displayed in the web interface are not unique since a gene can have several transcripts. These transcripts cannot be traced back to those determined using microarray chips or RNA sequencing, although they have been reported in the literature and collected during the first generation of the human metabolic reconstructions^17^. These transcript assignments have been retained in subsequent versions of the human metabolic reconstruction, although new genes have been added assuming only one transcript, specified as ‘.1’.

Each page contains additional information for each gene and external links to several resources, including Ensembl^76^, HGNC^77^, ChEMBL^78^, Uniprot^79^, Entrez Gene^80^, OMIM^81^ ^82^, Human Protein Atlas^83^, UCSC^84^, WikiGene^85^, and Gene Ontology^86^. Furthermore, connections with diseases and associated reactions are included similar to the other database entities.

### Microbe page

The microbe page provides details on the phylogeny, microbial metabolic reconstruction, external links to various databases and microbial resources. Additionally, each microbe has an associated set of numerical characteristics extracted from its reconstructions, reactions, metabolites, and gene contents along with a curated list of fermentation products and carbon sources. From each microbe page, the corresponding reconstruction in SBML and MAT formats as well as the genome in FASTA format can be downloaded.

### Food page

The food page provides information on the source where the nutritional information was extracted, a visual representation (i.e., pie chart) of its composition and the nutritional information separated by categories to facilitate the search for nutrients of interest.

### Diet page

The diet page contains information on each diet, a corresponding description and reference, as well as the composition of the diet in terms of included foodstuffs. From the diet page, each diet composition can be downloaded for use in metabolic modeling.

### Disease page

The disease page provides additional information on each disease, such as the type of disorder, inheritance pattern when applicable, phenotype and clinical symptoms, along with references and OMIM identifier. As with other entities in the VMH database, this detail page contains the connections with other entities in the database. Mutated genes and their associated reaction(s) are listed along with a biomarker profile linking metabolites.

### “ReconMaps” resource

The diagram editor CellDesigner (version 4.4)^21^ was used to manually draw the metabolic maps of the “ReconMaps” resource. Continuous quality control was achieved using a dedicated MATLAB (Mathworks, Inc.) code for map correction and manipulation. This code and the corresponding tutorial are freely available in the COBRA Toolbox^13^ (https://opencobra.github.io/cobratoolbox/).

From the ReconMaps resource, exchange reactions and reactions belonging to the subsystem “xenobiotic metabolism’ were excluded. Recon3D has nine compartments: extracellular [e], cytosol [c], mitochondria [m], mitochondrial intermembrane space [i], endoplasmic reticulum [r], lysosome [l], peroxisome [x], Golgi apparatus [g], and nucleus [n], which formed the basis of the organization of the map. Metabolites are represented as nodes identified by the metabolite name abbreviation and the compartment where the metabolite is present (e.g., water in the mitochondria = h2o[m]). For those compartments associated with cellular organelles, reaction and metabolite information is extracted and represented individually. Six submaps were generated, and all information for a specific organelle was compiled. For the mitochondria, mitochondrial [m] and mitochondrial intermembrane space [i] are represented in the same submap. Each submap is drawn by representing internal reactions in the center and transport reactions on the boundaries. Therefore, all metabolites transported between organelles and organelle/cytosol/extracellular compartments are located in the boundaries of the compartments. Combining the six submaps yielded ReconMap3.

We used the Interaction NEtwoRk visualization (MINERVA) web service^22^, to allow for content queries, custom dataset visualizations and feedback submission to manual curators. Custom datasets, particularly simulation results, can be visualized in a map context using the COBRA Toolbox^13^.

### Pre-defined diets and diet designer flux calculation

The 11 pre-defined diets available in the VMH database were designed by a nutrition professional following the caloric content based on the average recommended daily intake (i.e., 2500 calories for a male person). The diets consist of a one-day meal plan and include information about the energy content, fatty acids, amino acids, carbohydrates, dietary fibers, vitamins, minerals, and trace elements. The information for the nutritional composition or the foods and dishes has been provided by the “Österreichische Nährwerttabelle” (http://www.oenwt.at/content/naehrwert-suche/). The fluxes were calculated by converting the nutrient amount present in the foodstuff portions from grams to millimoles per human per day. The molecular mass of each metabolite is calculated. After converting the units, we determined the amount of that metabolite in the portion of food using the database’s nutritional information:

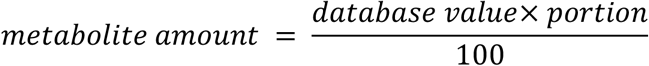

Then, we convert the metabolite amount into a flux value (mmol per person per day) using the following formula:

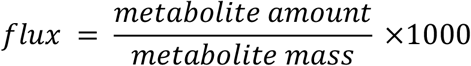

The fluxes are downloadable as text files, and each metabolite is associated with the corresponding exchange reactions using the VMH ID. These fluxes can be integrated with a metabolic model, e.g., Recon3D, using the COBRA Toolbox^13^. However, certain limitations are worth mentioning. Nutrient content from food labels and databases usually lacks several details that can influence the designed diets. While the USDA database for food references is exceptional in its detail, users must consider additional aspects. For instance, dietary fibers are counted as a unique nutrient. We have chosen not to map this metabolite group to any metabolite because several fibers can be consumed by different microbes, and depending on the selected food items, the user will have to make different assumptions.

### Data and software availability

The VMH database is freely available. The database content and search results are downloadable. Users can provide feedback through the different platforms of the website, and these issues will be curated and integrated into the database. The VMH database is freely accessible online at http://vmh.life.

### Interaction profiles of the microbes

A distance matrix between all the interaction profiles of the microbes was calculated using the Jaccard distance. A TSNE^38^ was performed on the distance matrix with 3000 iterations and a perplexity value of 30. For the comparison of the microbe consumption profile in Figure 4, the metabolites used are detailed in Supp. Table 2.

---

**Supp. Table 1:** Foodstuffs in VMH with the highest concentrations of fructose and galactose. Source of food nutritional information: US Department of Agriculture, Agricultural Research Service, Nutrient Data Laboratory. USDA National Nutrient Database for Standard Reference, Release 28. Version Current: September 2015.

| A – Fructose-rich foods |  | Values in g per 100 g |  |  |
| --- | --- | --- | --- | --- |
| Food | Producer | Fructose | Glucose | Total Sugar |
| Sweetener, syrup, agave |  | 55.6 | 12.43 | 68.03 |
| Agave, dried (Southwest) |  | 42.83 | 3.48 | 50.7 |
| Honey |  | 40.94 | 35.75 | 82.12 |
| Dates, medjool |  | 31.95 | 33.68 | 66.47 |
| Raisins, seedless |  | 29.68 | 27.75 | 59.19 |
| Cranberries, dried, sweetened |  | 26.96 | 29.69 | 72.56 |
| Figs, dried, uncooked |  | 22.93 | 24.79 | 47.92 |
| Lemonade-flavor drink, powder |  | 22.73 | 2.26 | 97.15 |
| Jujube, Chinese, fresh, dried |  | 20.62 | 18.28 | 0 |
| Lemonade, frozen concentrate, pink |  | 20.06 | 18.57 | 46.46 |
| Dates, deglet noor |  | 19.56 | 19.87 | 63.35 |
| Lemonade, frozen concentrate, white |  | 17.99 | 16.3 | 44.46 |
| Agave, cooked (Southwest) |  | 17.57 | 1.58 | 20.87 |
| Sauce, barbecue, SWEET BABY RAY'S, original | Sweet baby Ray's, Inc. | 17.52 | 20.85 | 38.37 |
| Beverages, lemonade, powder |  | 17.5 | 2.75 | 94.7 |
| Formulated bar, POWER BAR, chocolate |  | 15.96 | 11.94 | 30.07 |
| MCDONALD'S, Sweet 'N Sour Sauce | McDonald's Corporation | 15.63 | 18.76 | 35.79 |
| MCDONALD'S, Barbeque Sauce | McDonald's Corporation | 15.44 | 18.27 | 34.31 |
| Sauce, barbecue, KRAFT, original | Kraft Foods, Inc. | 14.58 | 16.65 | 32.26 |
| Sauce, barbecue |  | 14.17 | 16.39 | 33.24 |
| **B – Galactose-rich foods** |  | **Values in g per 100 g** |  |  |
| **Food** | **Producer** | **Galactose** | **Glucose** | **Total Sugar** |
| Formulated bar, SLIM-FAST OPTIMA meal bar, milk chocolate peanut | Slim-Fast Foods Company | 5.62 | 1.24 | 25 |
| Honey |  | 3.1 | 35.75 | 82.12 |
| Dulce de Leche |  | 1.03 | 1.7 | 49.74 |
| Celery, cooked, boiled, drained, without salt |  | 0.85 | 0.71 | 2.37 |
| Celery, cooked, boiled, drained, with salt |  | 0.85 | 0.71 | 2.37 |
| Beets, canned, regular pack, solids and liquids |  | 0.8 | 0.28 | 6.53 |
| Yogurt, Greek, nonfat, vanilla, CHOBANI | Chobani | 0.68 | 0.3 | 7.61 |
| Yogurt, Greek, vanilla, nonfat |  | 0.6 | 0.32 | 9.54 |
| Yogurt, Greek, vanilla, lowfat |  | 0.6 | 0.32 | 9.54 |
| Cherries, sweet, raw |  | 0.59 | 6.59 | 12.82 |
| Yogurt, Greek, nonfat, strawberry, DANNON OIKOS | Danone | 0.56 | 0.25 | 11.63 |
| Yogurt, Greek, nonfat, strawberry, CHOBANI | Chobani | 0.55 | 0.77 | 10.86 |
| Yogurt, Greek, strawberry, nonfat |  | 0.55 | 0.65 | 11.27 |
| Yogurt, Greek, strawberry, DANNON OIKOS | Danone | 0.54 | 0.3 | 11 |
| Yogurt, Greek, nonfat, vanilla, DANNON OIKOS | Danone | 0.54 | 0.27 | 11.4 |
| Yogurt, Greek, strawberry, lowfat |  | 0.53 | 0.54 | 11.23 |
| Celery, raw |  | 0.48 | 0.4 | 1.34 |
| T.G.I. FRIDAY'S, fried mozzarella | T.G.I Friday's | 0.4 | 0.5 | 1.45 |
| Spices, onion powder |  | 0.36 | 0.73 | 6.63 |
| Corn, sweet, white, canned, whole kernel, drained solids |  | 0.36 | 0.83 | 2.42 |

**Supp. Table 2:** Metabolites used in the polysaccharide consumption comparison of the microbe resource used in Figure 4.

| Polysaccharide | VMH Metabolite(s) identifiers |
| --- | --- |
| Mucus-O-Glycans | core2, core3, core4, core5, core6, core7, core8 |
| Pullulan | pullulan1200 |
| Glycogen | glygn1, glygn2, glygn3, glygn4, glygn5 |
| Amylopectin | amylopect900 |
| Inulin | inulin |
| Levan | levan1000 |
| Heparin | hspg |
| Hyaluronan | ha |
| Polygalacturonate | galur |
| Chondroitin sulfate | cspg_a, cspg_b, cspg_c |
| Rhamnogalacturonan | rhamnogalurI, rhamnogalurII |
| Pectic galactan | pecticgal |
| Arabinogalactan | arabinogal |
| Arabinan | arabinan101 |
| Oat spelt xylan | xylan |
| Galactomannan | glcmannan |
| Xyloglucan | xyluglc |
| Beta-glucan | bglc |
| Cellobiose | cellb |
| Laminarin | lmn30 |
| Lichenin | lichn |
| Dextran | dextran40 |
| Alpha-mannan | mannan |

## References

1. Rigden, D.J. & Fernandez, X.M. The 2018 Nucleic Acids Research database issue and the online molecular biology database collection. Nucleic Acids Res 46, D1–D7 (2018).

2. Swainston, N. et al. biochem4j: Integrated and extensible biochemical knowledge through graph databases. PLoS One 12, e0179130 (2017).

3. Palsson, B. Systems biology: properties of reconstructed networks. (Cambridge University Press, Cambridge; 2006).

4. Thiele, I. & Palsson, B.O. A protocol for generating a high-quality genome-scale metabolic reconstruction. Nature protocols 5, 93–121 (2010).

5. Brunk, E. et al. Recon3D enables a three-dimensional view of gene variation in human metabolism. Nature Biotechnology (2018).

6. Magnusdottir, S. et al. Generation of genome-scale metabolic reconstructions for 773 members of the human gut microbiota. Nat Biotechnol 35, 81–89 (2017).

7. Heinken, A. et al. Functional metabolic map of Faecalibacterium prausnitzii, a beneficial human gut microbe. Journal of bacteriology 196, 3289–3302 (2014).

8. Shoaie, S. et al. Understanding the interactions between bacteria in the human gut through metabolic modeling. Scientific reports 3, 2532 (2013).

9. Heinken, A. & Thiele, I. Systematic prediction of health-relevant human-microbial co-metabolism through a computational framework. Gut Microbes 6, 120–130 (2015).

10. Yizhak, K. et al. Phenotype-based cell-specific metabolic modeling reveals metabolic liabilities of cancer. eLife 3 (2014).

11. Bordbar, A. et al. Elucidating dynamic metabolic physiology through network integration of quantitative time-course metabolomics. Scientific reports 7, 46249 (2017).

12. Aurich, M.K., Fleming, R.M. & Thiele, I. MetaboTools: A Comprehensive Toolbox for Analysis of Genome-Scale Metabolic Models. Frontiers in physiology 7, 327 (2016).

13. Heirendt, L. et al. Creation and analysis of biochemical constraint-based models: the COBRA Toolbox v3.0. arXiv preprint (2017).

14. Nielsen, J. Systems Biology of Metabolism: A Driver for Developing Personalized and Precision Medicine. Cell metabolism 25, 572–579 (2017).

15. Zhang, C. & Hua, Q. Applications of Genome-Scale Metabolic Models in Biotechnology and Systems Medicine. Frontiers in physiology 6, 413 (2015).

16. Thiele, I. et al. A community-driven global reconstruction of human metabolism. Nat Biotechnol 31, 419–425 (2013).

17. Duarte, N.C. et al. Global reconstruction of the human metabolic network based on genomic and bibliomic data. Proceedings of the National Academy of Sciences of the United States of America 104, 1777–1782 (2007).

18. Swainston, N. et al. Recon 2.2: from reconstruction to model of human metabolism. Metabolomics: Official journal of the Metabolomic Society 12, 109 (2016).

19. Gonzalez, G.A.P. et al. Comparative evaluation of atom mapping algorithms for balanced metabolic reactions: application to Recon 3D. Journal of cheminformatics 9, 39(2017).

20. Noor, E., Haraldsdottir, H.S., Milo, R. & Fleming, R.M. Consistent estimation of Gibbs energy using component contributions. PLoS Comput Biol 9, e1003098 (2013).

21. Funahashi, A. et al. CellDesigner 3.5: A Versatile Modeling Tool for Biochemical Networks. Proceedings of the IEEE 96, 1254–1265 (2008).

22. Gawron, P. et al. MINERVA-a platform for visualization and curation of molecular interaction networks. NPJ Syst Biol Appl 2, 16020 (2016).

23. Noronha, A. et al. ReconMap: an interactive visualization of human metabolism. Bioinformatics (Oxford, England) 33, 605–607 (2017).

24. Wishart, D.S. et al. HMDB 3.0--The Human Metabolome Database in 2013. Nucleic Acids Res 41, D801–807 (2013).

25. Kanehisa, M., Goto, S., Sato, Y., Furumichi, M. & Tanabe, M. KEGG for integration and interpretation of large-scale molecular data sets. Nucleic Acids Res 40, D109–114 (2012).

26. Fujita, K.A. et al. Integrating pathways of Parkinson’s disease in a molecular interaction map. Molecular neurobiology 49, 88–102 (2014).

27. Kuperstein, I. et al. Atlas of Cancer Signalling Network: a systems biology resource for integrative analysis of cancer data with Google Maps. Oncogenesis 4, e160 (2015).

28. Sompairac, N. et al. Metabolic and signalling network map integration: application to cross-talk studies and omics data analysis in cancer. BioRxiv preprint (Submitted).

29. Ostaszewski, M. et al. Community-driven roadmap for integrated disease maps. Brief Bioinform (2018).

30. Qin, J. et al. A human gut microbial gene catalogue established by metagenomic sequencing. Nature 464, 59–65 (2010).

31. Sahoo, S., Franzson, L., Jonsson, J.J. & Thiele, I. A compendium of inborn errors of metabolism mapped onto the human metabolic network. Molecular bioSystems 8, 2545–2558 (2012).

32. Rahman, J., Noronha, A., Thiele, I. & Rahman, S. Leigh map: A novel computational diagnostic resource for mitochondrial disease. Ann Neurol 81, 9–16 (2017).

33. Köhler, S. et al. The Human Phenotype Ontology project: linking molecular biology and disease through phenotype data. Nucleic Acids Res 42, D966-D974 (2014).

34. U.S. Department of Agriculture, A.R.S. (2011).

35. Elmadfa, I. Österreichischer Ernährungsbericht 2012, Edn. 1. (Vienna; 2012).

36. Thiele, I. et al. Personalized whole-body models integrate metabolism, physiology, and the gut microbiome. https://www.biorxiv.org/content/early/2018/01/29/255885 (Submitted).

37. Round, J.L. & Mazmanian, S.K. The gut microbiota shapes intestinal immune responses during health and disease. Nature reviews 9, 313–323 (2009).

38. Maaten, L.v.d. & Hinton, G. Visualizing data using t-SNE. Journal of Machine Learning Research 9, 2579–2605 (2008).

39. Ndeh, D. & Gilbert, H.J. Biochemistry of complex glycan depolymerisation by the human gut microbiota. FEMS microbiology reviews 42, 146–164 (2018).

40. Desai, M.S. et al. A Dietary Fiber-Deprived Gut Microbiota Degrades the Colonic Mucus Barrier and Enhances Pathogen Susceptibility. Cell 167, 1339–1353. e1321 (2016).

41. Becker, N., Kunath, J., Loh, G. & Blaut, M. Human intestinal microbiota: characterization of a simplified and stable gnotobiotic rat model. Gut Microbes 2, 25–33 (2011).

42. Sanchez, R. & Kauffman, F. Regulation of Xenobiotic Metabolism in the Liver. Comprehensive Toxicology 9, 109–128 (2010).

43. Howell, S.R., Hazelton, G.A. & Klaassen, C.D. Depletion of hepatic UDP-glucuronic acid by drugs that are glucuronidated. J Pharmacol Exp Ther 236, 610–614 (1986).

44. Sahoo, S., Haraldsdottir, H., Fleming, R.M. & Thiele, I. Modeling the effects of commonly used drugs on human metabolism. FEBS Journal 282, 297–317 (2015).

45. Bánhegyi, G., Garzó, T., Antoni, F. & Mandl, J. Glycogenolysis-and not gluconeogenesis-is the source of UDP-glucuronic acid for glucuronidation. Biochimica et Biophysica Acta (BBA)-General Subjects 967, 429–435 (1988).

46. Conlee, R.K., Lawler, R.M. & Ross, P.E. Effects of glucose or fructose feeding on glycogen repletion in muscle and liver after exercise or fasting. Annals of nutrition and metabolism 31, 126–132 (1987).

47. Décombaz, J. et al. Fructose and galactose enhance postexercise human liver glycogen synthesis. Medicine and science in sports and exercise 43, 1964–1971 (2011).

48. Nickerson, K.P. & McDonald, C. Crohn’s disease-associated adherent-invasive Escherichia coli adhesion is enhanced by exposure to the ubiquitous dietary polysaccharide maltodextrin. PLoS One 7, e52132 (2012).

49. Nickerson, K.P., Chanin, R. & McDonald, C. Deregulation of intestinal anti-microbial defense by the dietary additive, maltodextrin. Gut microbes 6, 78–83 (2015).

50. Truswell, A.S., Seach, J.M. & Thorburn, A. Incomplete absorption of pure fructose in healthy subjects and the facilitating effect of glucose. The American journal of clinical nutrition 48, 1424–1430 (1988).

51. Stringer, A.M. et al. Chemotherapy-induced mucositis: the role of gastrointestinal microflora and mucins in the luminal environment. The journal of supportive oncology 5, 259–267 (2007).

52. Takakura, A. et al. Rapid deconjugation of SN-38 glucuronide and adsorption of released free SN-38 by intestinal microorganisms in rat. Oncol Lett 3, 520–524 (2012).

53. Nyhan, W.L. Disorders of purine and pyrimidine metabolism. Molecular genetics and metabolism 86, 25–33 (2005).

54. Wurtman, R.J., Regan, M., Ulus, I. & Yu, L. Effect of oral CDP-choline on plasma choline and uridine levels in humans. Biochemical pharmacology 60, 989–992 (2000).

55. Simmonds, H., Webster, D., Becroft, D. & Potter, C. Purine and pyrimidine metabolism in hereditary orotic aciduria: some unexpected effects of allopurinol. European journal of clinical investigation 10, 333–339 (1980).

56. Heinken, A., Sahoo, S., Fleming, R.M. & Thiele, I. Systems-level characterization of a host-microbe metabolic symbiosis in the mammalian gut. Gut Microbes 4, 28–40 (2013).

57. Baumgartner, M.R. et al. Proposed guidelines for the diagnosis and management of methylmalonic and propionic acidemia. Orphanet J Rare Dis 9, 130 (2014).

58. Durrer, K.E., Allen, M.S. & von Herbing, I.H. Genetically engineered probiotic for the treatment of phenylketonuria (PKU); assessment of a novel treatment in vitro and in the PAHenu2 mouse model of PKU. PloS one 12, e0176286 (2017).

59. Kotiranta, A., Lounatmaa, K. & Haapasalo, M. Epidemiology and pathogenesis of Bacillus cereus infections. Microbes and infection 2, 189–198 (2000).

60. Mollstam, B. & Connolly, E. (Google Patents, 2005).

61. Berry, D., Kaplan, J. & Rahman, S. (Google Patents, 2017).

62. Nilsson, A., Mardinoglu, A. & Nielsen, J. Predicting growth of the healthy infant using a genome scale metabolic model. NPJ Syst Biol Appl 3, 3 (2017).

63. Heinken, A. et al. Personalized modeling of the human gut microbiome reveals distinct bile acid deconjugation and biotransformation potential in healthy and IBD individuals. BioRxiv preprint (2017).

64. Opdam, S. et al. A Systematic Evaluation of Methods for Tailoring Genome-Scale Metabolic Models. Cell systems 4, 318–329 e316 (2017).

65. Claesson, M.J. et al. Gut microbiota composition correlates with diet and health in the elderly. Nature 488, 178–184 (2012).

66. Zeevi, D. et al. Personalized Nutrition by Prediction of Glycemic Responses. Cell 163, 1079–1094 (2015).

67. Coles, S.J., Day, N.E., Murray-Rust, P., Rzepa, H.S. & Zhang, Y. Enhancement of the chemical semantic web through the use of InChI identifiers. Organic & biomolecular chemistry 3, 1832–1834 (2005).

68. Weininger, D. SMILES, a chemical language and information system. 1. Introduction to methodology and encoding rules. Journal of Chemical Information and Computer Sciences 28, 31–36 (1988).

69. Shin, S.-Y. et al. An atlas of genetic influences on human blood metabolites. Nature genetics 46, 543–550 (2014).

70. Engelke, U. et al. Handbook of 1H-NMR spectroscopy in inborn errors of metabolism: body fluid NMR spectroscopy and in vivo MR spectroscopy. Heilbronn: SPS Verlagsgesellschaft (2007).

71. Chen, R. et al. Personal omics profiling reveals dynamic molecular and medical phenotypes. Cell 148, 1293–1307 (2012).

72. Illig, T. et al. A genome-wide perspective of genetic variation in human metabolism. Nature genetics 42, 137–141 (2010).

73. Holle, R., Happich, M., Löwel, H., Wichmann, H. & group, M.K.s. KORA-a research platform for population based health research. Das Gesundheitswesen 67, 19–25 (2005).

74. Mittelstrass, K. et al. Discovery of sexual dimorphisms in metabolic and genetic biomarkers. PLoS genetics 7, e1002215 (2011).

75. Tatusov, R.L., Galperin, M.Y., Natale, D.A. & Koonin, E.V. The COG database: a tool for genomescale analysis of protein functions and evolution. Nucleic Acids Res 28, 33–36 (2000).

76. Cunningham, F. et al. Ensembl 2015. Nucleic Acids Res 43, D662–D669(2014).

77. Gray, K.A., Yates, B., Seal, R.L., Wright, M.W. & Bruford, E.A. Genenames. org: the HGNC resources in 2015. Nucleic Acids Res, gku1071 (2014).

78. Bento, A.P. et al. The ChEMBL bioactivity database: an update. Nucleic Acids Res 42, D1083-D1090 (2014).

79. Consortium, U. UniProt: a hub for protein information. Nucleic Acids Res, gku989 (2014).

80. Maglott, D., Ostell, J., Pruitt, K.D. & Tatusova, T. Entrez Gene: gene-centered information at NCBI. Nucleic Acids Res 39, D52–D57 (2010).

81. Amberger, J., Bocchini, C.A., Scott, A.F. & Hamosh, A. McKusick’s online Mendelian inheritance in man (OMIM®). Nucleic Acids Res 37, D793–D796 (2008).

82. Hamosh, A., Scott, A.F., Amberger, J.S., Bocchini, C.A. & McKusick, V.A. Online Mendelian Inheritance in Man (OMIM), a knowledgebase of human genes and genetic disorders. Nucleic Acids Res 33, D514–D517 (2005).

83. Uhlen, M. et al. Towards a knowledge-based human protein atlas. Nature biotechnology 28, 1248–1250 (2010).

84. Tyner, C. et al. The UCSC Genome Browser database: 2017 update. Nucleic Acids Res 45, D626–D634 (2016).

85. Hoffmann, R. A wiki for the life sciences where authorship matters. Nat Genet 40, 1047–1051 (2008).

86. Consortium, G.O. The Gene Ontology (GO) database and informatics resource. Nucleic Acids Res 32, D258–D261 (2004).

